# Applying Unet for extraction of vascular metrics from T1-weighted and T2-weighted MRI

**DOI:** 10.1101/2022.12.18.520922

**Authors:** Farnaz Orooji, Russell Butler

## Abstract

We apply deep learning to the problem of segmenting the arterial system from T1w and T2w images. We use the freely available 7-Tesla ‘forrest’ dataset from OpenNeuro, (which contains TOF, T1w, and T2w) and use supervised learning with T1w or T2w as input, and TOF segmentation as ground truth, to train a Unet architecture capable of segmenting arteries and quantifying arterial diameters from T1w or T2w images alone. We demonstrate arterial segmentations from both T1w and T2w images, and show that T2w images have sufficient vessel contrast to estimate arterial diameters comparable to those estimated from TOF. We then apply our Unet to T2w images from a separate dataset (IXI) and show our model generalizes to images acquired at different field strength. We consider this work proof-of-concept that arterial segmentations can be derived from MRI sequences with poor contrast between arteries and surrounding tissue (T1w and T2w), due to the ability of deep convolutional networks to extract complex features based on local image intensity. Future work will focus on improving the generalizability of the network to non-forrest datasets, with the eventual goal of leveraging the entire pre-existing corpus of neuroimaging data for study of human cerebrovasculature.

## 1 Introduction

### 1.1 Cerebrovasculature

The human brain consumes about 20% of daily intake of oxygen and glucose through blood supplied to the cortex via arteries, arterioles, and capillaries, and drained via the venous system and large sinuses. Arteries, which carry oxygenated blood towards the brain, are thought to play a key role in several neurodegenerative diseases (Sweeney et al., 2018) including Alzheimer’s and Multiple Sclerosis, and are critical in planning arterial-spin-labeling (ASL) experiments (Hartkamp et al., 2013). The malfunction of this system can lead to disease or death. The structure of cerebral vessels is complex, and its 3D representation is an enormous step forward in improving diagnostic accuracy and medical education (Akkus et al., 2017). The development of diagnostic imaging techniques, such as Magnetic Resonance Angiography (MRA) and Computed Tomography Angiography (CTA), enable to generate volumetric angiographic data whose post-processing provide a 3D reconstruction of vascular networks. Despite the existence of many neuroimaging tools for segmenting and quantifying the gray and white matter of the brain, few publicly available methods exist for non-invasive investigation of cerebral vasculature (Butler et al., 2017, 2019, 2020; Cote et al., 2021; Forouhandehpour et al., 2021; Jiang et al., 2022).

### 1.2 Deep learning for vascular segmentation

Deep learning methods have had good success recently in medical image analysis. In Livne et al. (2019), they performed arterial segmentation on 3T PEGASUS and 7T 7UP dataset with Unet model and half Unet model. In half Unet the number of channels is reduced by half in each layer. They used three metrics: Dice coefficient, 95% Hausdorff distance (95HD) and average Hausdorff distance (AVD), and showed that the results have a good performance on large vessels and sufficient performance for small vessels, based on these validation metrics and compared to the traditional segmentation graph-cut method. Phellan et al. (2017) proposed a fully convolutional network consisting of two convolutional layers followed by two fully connected layers, applied on TOF-MRA of five healthy subjects. These subjects were manually segmented and the final segmentation results were compared with the ground truth by Dice similarity coefficient, showing promising results in automatic vascular segmentation. Xiancheng et al. (2018) presents a training strategy for a Unet that relies on data augmentation. The objective of the model is to get evidence about a variety of eye diseases by using the shape, size, and arteriovenous crossing types. The parameters of the model such as the number of layers and their configuration were selected via experimentation and were balanced between the training time and the accuracy of the model. Dropout layers were also introduced between the convolutional layers to improve the training performance. The advantages of the proposed model are downscaling the network and reducing the number of feature vectors at each layer. Thus, the model obtains results equivalent to the original Unet, but the training became easier, and its time was significantly reduced.

The other study (de Vos et al., 2021) examined vessel segmentation of TOF-MRA on patch training on both 2D and 3D Unet. They examined the effect of data augmentation such as Gaussian blur, rotation and flipping on images from 3-Tesla dataset. For 2D and 3D images they also used different patch sizes. They showed that when the model was trained with whole slices, superior results were achieved based on three metrics (dice similarity coefficient, modified hausdorff distance and volumetric similarity).

Alom et al. (2018) propose two models for segmentation tasks; A RCNN and a Recurrent Residual Convolutional Neural Network (RRCNN), both based on Unet models. The models were evaluated on three datasets (blood vessel segmentation in the retina images, skin cancer segmentation, and lung lesion segmentation). Their results show that 1) residual units help when training deep architectures, 2) feature accumulation with recurrent residual convolutional layers ensures better feature representation for segmentation tasks and 3) it is possible to design a better Unet architecture with the same number of network parameters with better performance for medical image segmentation. The main problem with the above methods is that TOF images are not acquired in standard neuroimaging protocols, instead, T1-weighted (T1w) or T2-weighted (T2w) images are acquired to obtain information about the brain’s structure such as gray matter boundaries. Therefore, we have a large, preexisting corpus of neuroimaging data but we are limited in our ability to quantify the arterial system from this corpus due to the fact that segmenting arteries from T1w or T2w images is not straightforward. To address this we create a ground truth by registering single subject TOF to single subject T1 and T2, in order to train a deep neural network capable of segmenting brain arteries directly from T1 or T2 images.

## 2 Materials and Methods

### 2.1 Dataset

In this study, we used the OpenNeuro’s ‘Forrest’ dataset which contains 20 subjects (Gorgolewski et al., 2017). From Forrest dataset, we downloaded full brain volumetric images, for T1 and T2 images we have [274 × 384 × 384] slices in NIFTI format, and TOF images with [480 × 640 × 163] slices in NIFTI format recorded with a Siemens MR scanner at 7-Tesla using a 3D multi-slab sequence (0.3 mm isotropic voxel size). Both T1/T2 images have the resolution 0.7 × 0.6 × 0.6 mm.

### 2.2 Preparing Input for Unet model

Due to the multi-modal nature of our dataset, we must use image registration to put all images into the same space. Here, the target image is the TOF, because it has a limited field of view and a higher spatial resolution than the T1 or T2 images.

#### 2.2.1 Image Registration in T1-weighted and T2-weighted image

The best registration between TOF and T1 could be found by first registering the T2 to the TOF using a 6 degree of freedom (DOF) linear transform by FLIRT FSL and then a second fine-tuning registration using 9 DOF (from T2 in TOF to TOF again). This second registration input and reference were rough segmentation of arteries in T2 and TOF, respectively. We then registered the T1 to the T2 and applied the above two matrices to bring T1 into TOF space. Figure 2.1, clearly shows the necessity of fine-tuning registration. In this figure, (A), we can see the first registration T2 in TOF space in axial view, and in the right column, shows the second finetuning registration T2 in TOF to TOF, also, it illustrates that the overlaid T2 and TOF perfectly aligned.

**Fig. 2.1.**
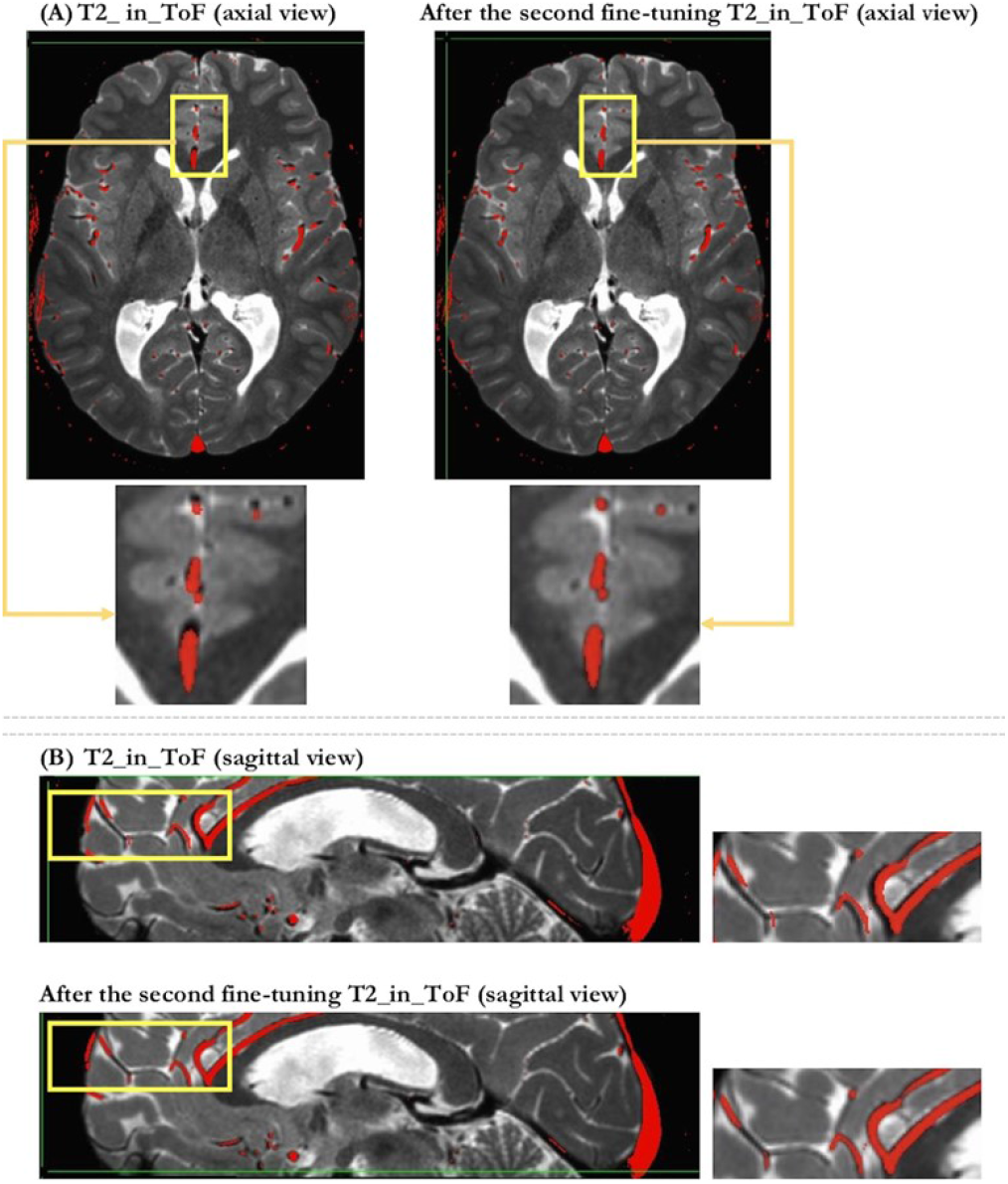
Image Registration

### 2.3 Ground truth generation

OpenNeuro’s Forrest Gump dataset does not contain expert delineations of the brain vasculature, however, the TOF images are of sufficient SNR to allow for a semi-automatic ground truth segmentation generation by applying the following steps.

1. Skull-stripped of T2-weighted image. skull strip is the process that isolate the brain from non-brain tissue and lead to get a better segmentation of the brain tissue (Kalavathi and Prasath, 2016). We apply Brain Extraction Tool - ‘BET’ (Cox, 1996) command to get the skull stripped T2w. Figure 2.2 (b)
2. Get the rough T2 in TOF space registration using 6 degree of freedom linear transform, with the ‘flirt’ command. Figure 2.2 (c)
3. Erode the T2 skull strip mask with the command ‘3dLocalstat’, to process data from a spatial neighborhood of each voxel, here we compute the maximum over cube (5,5,5) placed around each voxel. The output of this step is the eroded T2 mask. Figure 2.2 (d)
4. Bring the eroded T2 skull stripped mask into TOF space by applying the matrix that was brought from T2 in TOF space. Figure 2.2 (e)
5. Threshold the TOF image by choosing the image specific intensity percentage at which the threshold was determined (Wang et al., 2015). For each subject we adjusted the threshold value individually, because it may be different from subject to subject. After thresholding the TOF we multiply it by eroded skull stripped mask to remove the edge of the brain, because in the segmentation task we have to consider only the vessels inside the brain. This step is done by ‘3dcalc’ command, and the output of this step is TOF segmented vessels, Figure 2.2 (f). Figure 2.3, illustrates the obtained segmented vessels of one subject of our dataset with the maximum intensity projection-MIP (Sun and Parker, 1999).

**Fig. 2.2.**
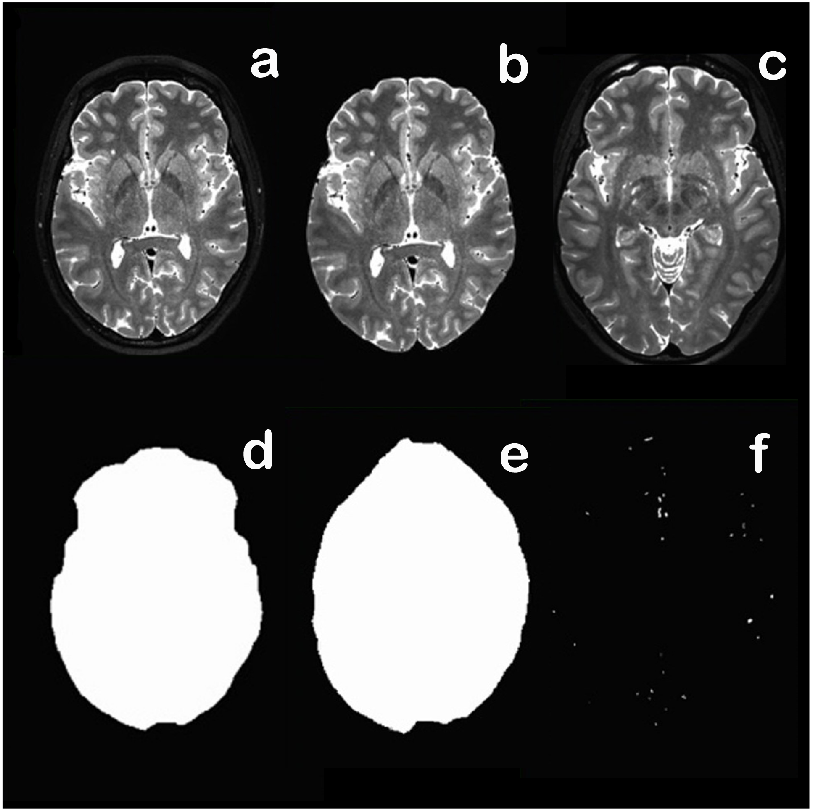
The steps of ground truth generation. (a) raw T2, (b) BET T2, (c) T2-in-TOF, (d) erode T2-mask, (e) erode TOF-mask and (f) TOF-vessels

**Fig. 2.3.**
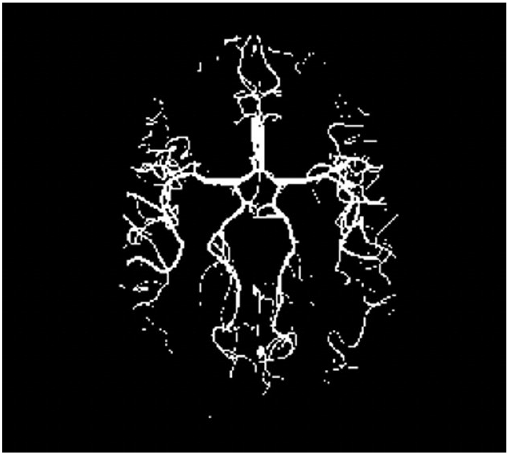
ground truth – the maximum intensity projection of ground truth-TOF

### 2.4 Implementation method

In this study, an arterial segmentation method based on Unet architecture, is proposed to address the problem of insufficient availability of some modalities such as TOF-MRA in medical research and diagnostic. Moreover, we compare the arterial segmentation results of Unet model with another deep learning segmentation model (Linknet) (Chaurasia and Culurciello, 2017).

The goal is to segmentation of the arteries from T1-weighted and T2-weighted images, to 1) see if these images contain sufficient detail for arterial segmentation and 2) determine which MRI sequence (T1 or T2) has better vessel information. To verify our results, we applied evaluation metrics and extracted vessel diameters in both TOF and synthesized-TOF. Also, we compare the result of Unet model with the Linknet. We train the Unet and Linknet model with T1 and T2 separately, according the Table 2.1, then comparing the segmentation results for both models. All T1w and T2w were registered in the TOF space.

**Table 2.1.** Summary of methods for training the Unet model

|  | Input | Labels |
| --- | --- | --- |
| 1 | T1-weighted image | segmented-TOF |
| 2 | T2-weighted image | segmented-TOF |

We implemented our network in Keras. The model was optimized using Adam optimizer plus Sigmoid activation since we perform binary segmentation. We carried out experiments on axial slices. The model was trained by feeding all 2D images, from whole dataset, each image is an axial T1-weighted or T2-weighted slice obtained from subjects of our dataset and corresponding segmented TOF which we have obtained as describe in section 2.3. The parameters were fine-tuned based on visual inspection while training data. All model training was performed on 2 Intel®Xeon CPU ®2.00 GHz @with Tesla V100-SXM2-16GB.

### 2.5 Evaluation metrics

We used two evaluation metrics to compare the performance of the Unet and Linknet architecture. IOU is computed based on the confusion matrix between prediction and ground truth by calculating the number of true positives (*TP*), true negatives (*TN*), false positives (*FP*), and false-negatives (*FN*). These variables are used to measure the performance of the network in terms of intersection over union (*IoU*), F1-Score. In terms of *IoU*, it is defined as,

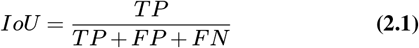

F1-Score is the weighted average of precision and recall, where an F-score reaches its best value at 1 and worst score at 0, precision, recall and F1-score are expressed as:

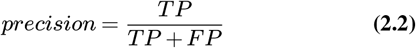

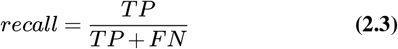

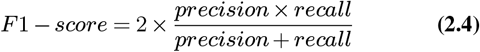

### 2.6 Skeletonization for diameter extraction

Another metric for quantification of the model, is to apply skeletonization to extract arterial diameters. We applied “kimimaro” skeletonize method (Silversmith and Bae., 2020). This algorithm works by finding a root point on a 3D object and then serially tracing paths via Dijkstra algorithm. We limited the analysis of the image by manually selecting the slices of both raw TOF and synthesized TOF from the Unet and Linknet. After applying skeletonization, by finding the vertices of segmented vessels in both images, we then could get the diameters of the segmented vessels in each vertex, finally we were able to compare diameters and found the correlations between these diameters. We have shown the correlation of skeletonization results in the next chapter.

## 3 Experimental Results

### 3.1 Segmentation results

Figure 3.1, illustrates the segmented arteries of Unet and Linknet model. Top row shows the 2D results of segmented arteries of T1 image, and the second row shows the results of segmented arteries of T2 image.

**Fig. 3.1.**
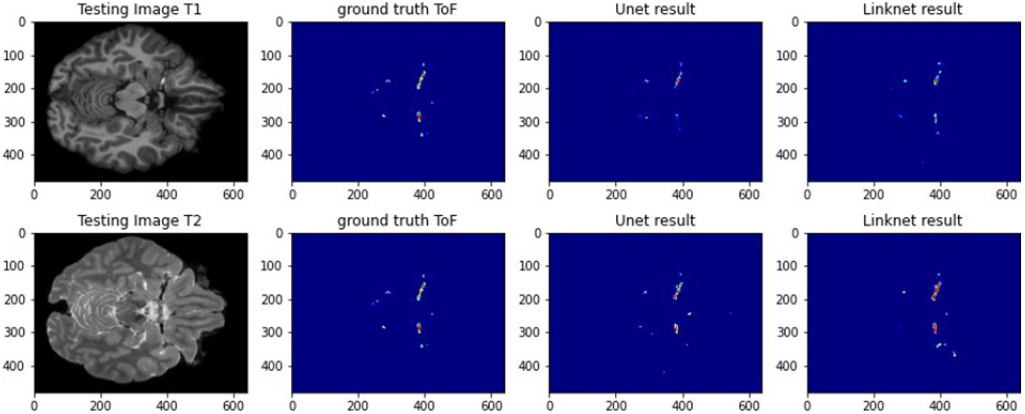
Comparison between Unet and Linknet on T1/T2

Figure 3.2 represents the corresponding maximum intensity projection of the result of segmented arteries Unet model, when the model was trained with T1 image (a). The next image represents the segmented arteries when the Linknet model trained with T1 image (b), and the top-right picture, shows the ground truth of test subject (c). The second row of images displays the same picture as above but zoomed in allowing for better visual evaluation of the results (d, e, and f).

**Fig. 3.2.**
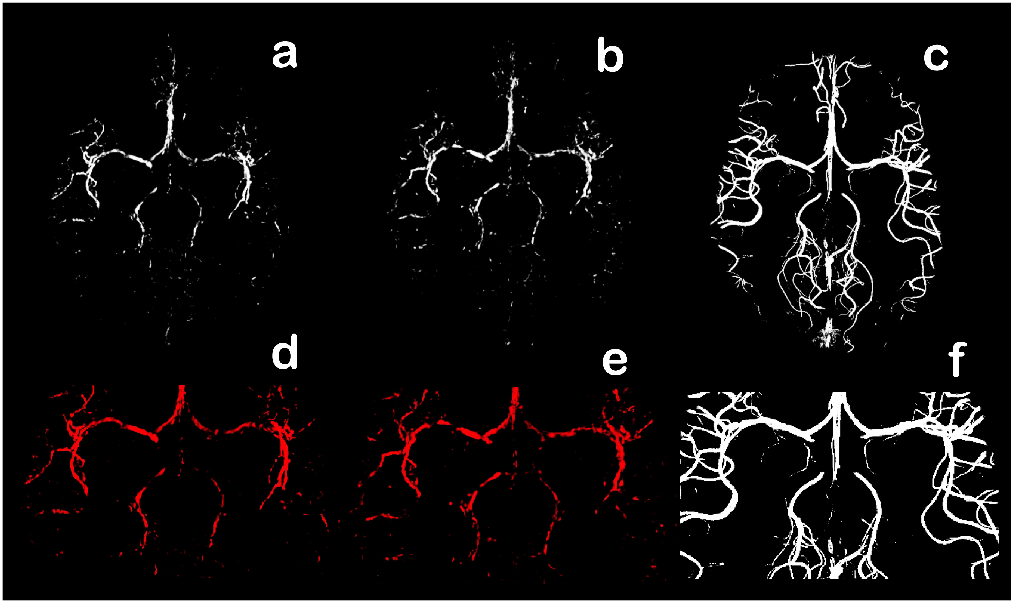
T1-weighted image prediction results. (a, c) Unet prediction, (b, d) Linknet prediction, (c, f) ground truth

Figure 3.3, illustrates the corresponding maximum intensity projection of the result of segmented arteries Unet model, when the model was trained with T2 image (a). The next picture represents the segmented arteries when the Linknet model trained using T1 image as input (b), and the top-right picture shows the ground truth of test subject. The next row show zoomed versions of the above pictures (d, e, and f). By visual evaluation, from the above figures, it can be concluded that T2 image gives more accurate arterial segmentation results when compare to the ground truth, as expected from the results of the Table 3.1.

**Table 3.1.** The evaluation metrics results of Unet/Linknet methods on both T1 and T2 image

| Method | Training on | F1-score | IoU-Score | Trainable parameters |
| --- | --- | --- | --- | --- |
| Unet | T1-weighted image | 0.82 | 0.68 | 24M |
| Linknet | T1-weighted image | 0.77 | 0.65 | 21M |
| **Unet** | **T2-weighted image** | **0.87** | **0.72** | 24M |
| Linknet | T2-weighted image | 0.81 | 0.69 | 21M |

Table 3.1 shows that the Unet model outperformed the Linknet approach also, in terms of the modality, T2-weighted image has better

**Fig. 3.3.**
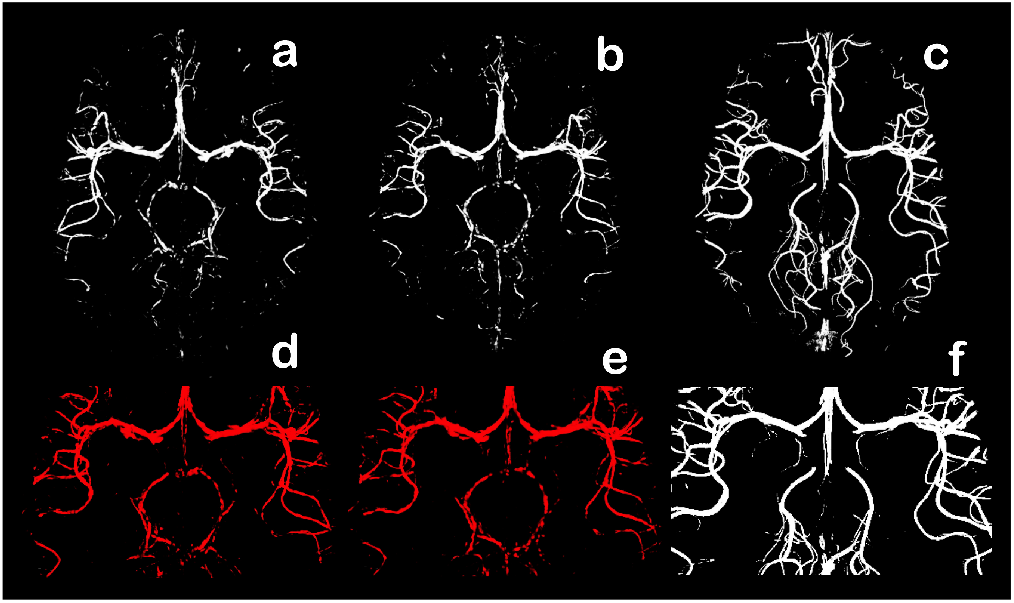
T2-weighted image prediction results. (a, c) Unet prediction, (b, d) Linknet prediction, (c, f) ground truth

F1-score and IOU-score 0.87, 0.72, respectively than T1-weighted.

### 3.2 Diameter extraction and correlation

Figure 3.4 illustrates the comparison between two models regarding the correlation map of diameters of the predicted segmentation vs the ground truth segmentation, using kimimaro skeletonization method to extract diameters. As expected from table 3.1, there is high positive correlation (0.93) with the Unet model trained using T2 images. In contrast, T1 image does not bring a high correlation between real and synthesized arterial diameters. Moreover, Unet model generalizes better than Linknet architecture.

**Fig. 3.4.**
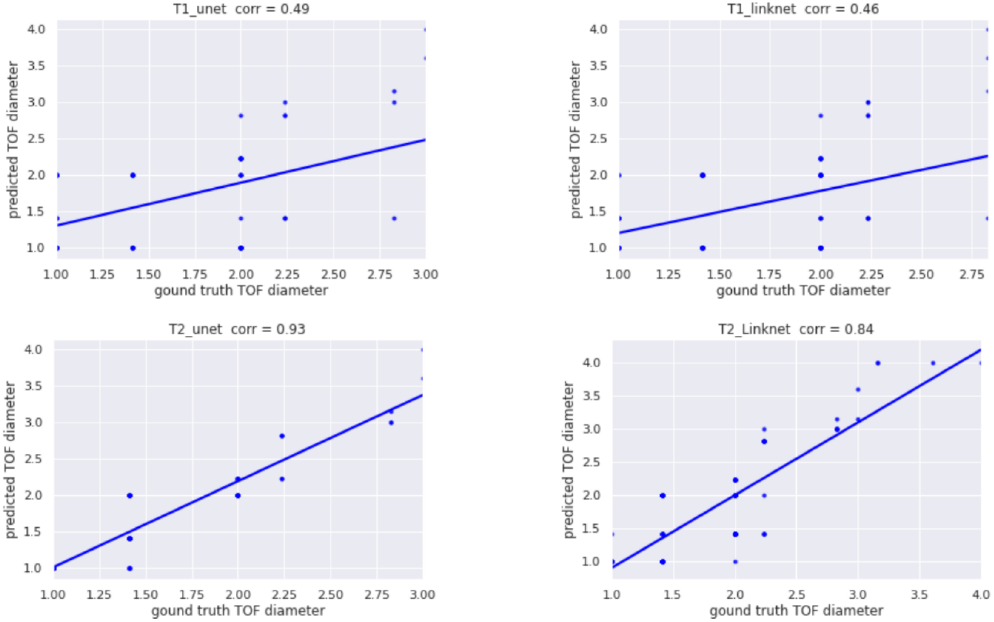
Correlation map of the diameters of the synthesized TOF vs the ground truth TOF

### 3.3 Experiment with combining T1 and T2

We performed additional experiments by training the models on both T1 and T2 images combined into a single training set. The aim was to experiment and compare the predicted segmentation task with the previous work. At this stage, according to the Table 3.2, the Unet model still has better evaluation results and T2-weighted image proves more accurate, in terms of the IOU-score and F1-score. Some vessels are missed when we combine T1 and T2 images with comparison when we train model with T2 image only.

**Table 3.2.** The evaluation metrics results of Unet/Linknet methods on both T2 and T1+T2 image

| Method | Training on | F1-score | IoU-Score | Trainable parameters |
| --- | --- | --- | --- | --- |
| **Unet** | **T2-weighted image** | **0.87** | **0.72** | 24M |
| Linknet | T2-weighted image | 0.81 | 0.69 | 21M |
| Unet | T1-weighted + T2-weighted image | 0.72 | 0.65 | 24M |
| Linknet | T1-weighted + T2-weighted image | 0.69 | 0.61 | 21M |

### 3.4 Testing the model on a separate dataset (IXI)

We chose our best model, Unet, trained with T2-weighted images from forrest dataset, and performed vessel segmentation on T2-weighted images from IXI dataset. This dataset contains 600 MR images from normal, healthy subjects. All the images are in the NIFTI format. The MR image acquisition protocol for each subject includes T1, T2, and TOF MRA images, the result from prediction on a single subject of IXI is below:

**Fig. 3.5.**
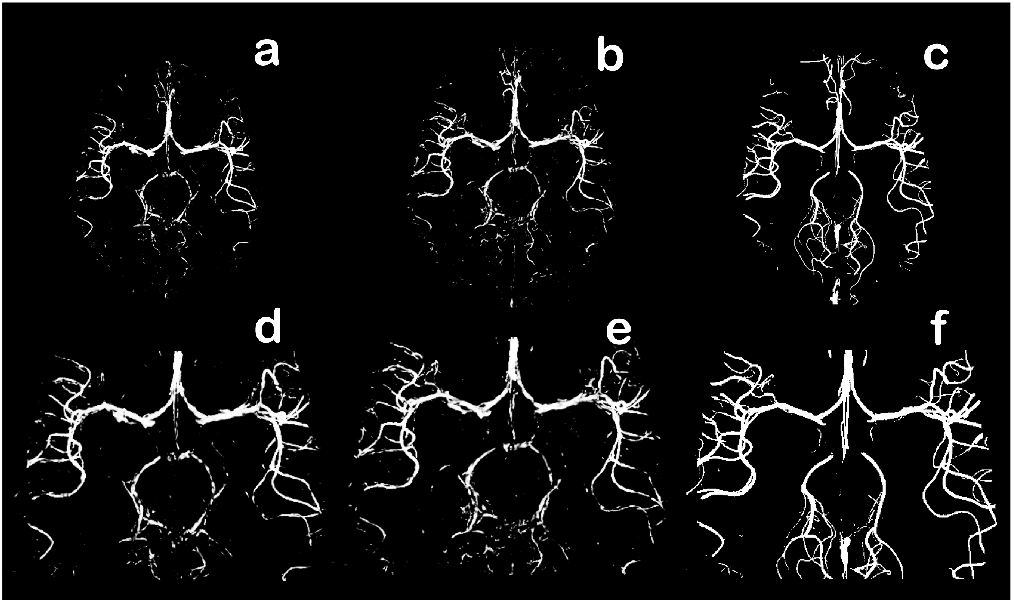
T2-weighted image prediction results. (a, d) Unet prediction with T2 image, (b, e) Unet prediction with T1+T2 images and (c, f) ground truth

**Fig. 3.6.**
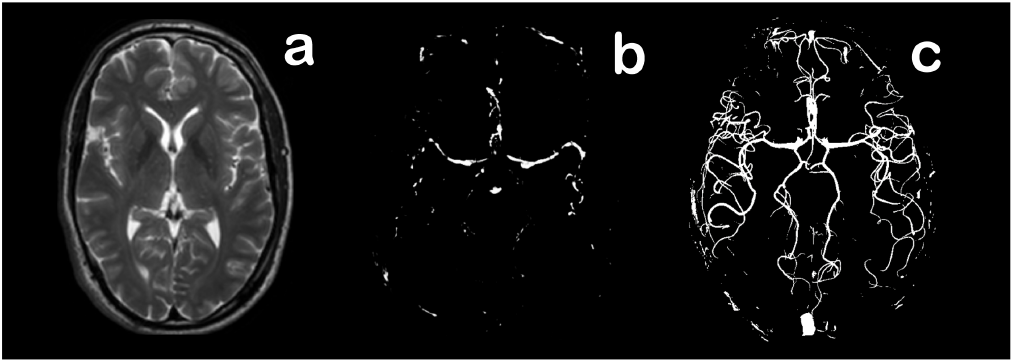
Prediction with IXI dataset. (a) T2 image, (b) TOF prediction with Unet and (c) Ground Truth

As can be seen above, the model was able to make some generalization to the T2-weighted image of the IXI dataset, but only the major arteries (middle cerebral arteries) were detected using our model. Further work is needed to improve the generalization capacity of our model, to achieve our end-goal of segmenting vasculature from a large pre-existing corpus of T1-weighted or T2-weighted images.

## 4 Discussions

Using 2D Unet architecture, we show it is possible to extract measures of vascular diameter from T2-weighted and T1-weighted images. To the best of our knowledge, this is the first study to examine the feasibility of extracting vascular segmentation and diameter metrics from T1-weighted or T2-weighted images. The fact that arterial diameter can be extracted from T1-weighted and T2-weighted images is unsurprising, because arteries are clearly visible on both T1 and T2. However, these images lack good contrast between arteries and surrounding tissue, and the details of the arteries in a T1 are different from arteries in TOF. Arteries appear brightly on the TOF in all brain areas, while in the T1, arteries are brighter in the brainstem/neck area, but darker and less prominent in more superior brain regions. Arteries are also visible on T2-weighted images as well as veins. This allows for synthesis of arterial trees in preexisting datasets, to perform additional analysis regarding vascular health.

We concluded that arteries were better segmented from the T2-weighted image than the T1 as we have shown the Figure 4.3. T2 has better vessel contrast than T1. This might be due to the fact that T2w image contrast contains some susceptibility effects due to T2* relaxation, increasing the contrast of arteries on T2w images relative to surrounding tissue.

We also experimented with another deep learning architecture, Linknet which was applied in the segmentation tasks. From the results that we have obtained (Table 3.1), we concluded that Unet model out performed this model. We should note that the 2D Unet has limitations, here we show axial slices (the model was trained on axial slices) but in the future, we plan to move to 3D Unet to obtain more accurate synthesis results. Also, our model was validated on two datasets only (IXI and forrest), and further work is necessary to verify how the model will perform on T2-weighted images acquired in different settings.

## References

Akkus, Z., Galimzianova, A., Hoogi, A., Rubin, D. L., and Erickson, B. J. (2017). Deep learning for brain MRI segmentation: State of the art and future directions. J. Digit. Imaging, 30(4):449–459.

Alom, M. Z., Hasan, M., Yakopcic, C., Taha, T. M., and Asari, V. K. (2018). Recurrent residual convolutional neural network based on U-Net (R2U-Net) for medical image segmentation. arXiv.

Butler, R., Bernier, P.-M., Lefebvre, J., Gilbert, G., and Whittingstall, K. (2017). Decorrelated input dissociates narrow band γ power and bold in human visual cortex. Journal of Neuroscience, 37(22):5408–5418.

Butler, R., Bernier, P.-M., Mierzwinski, G. W., Descoteaux, M., Gilbert, G., and Whittingstall, K. (2019). Cortical distance, not cancellation, dominates inter-subject EEG gamma rhythm amplitude. Neuroimage, 192:156–165.

Butler, R., Mierzwinski, G. W., Bernier, P. M., Descoteaux, M., Gilbert, G., and Whittingstall, K. (2020). Neurophysiological basis of contrast dependent BOLD orientation tuning. Neuroimage, 206(116323):116323.

Chaurasia, A. and Culurciello, E. (2017). LinkNet: Exploiting encoder representations for efficient semantic segmentation. In 2017 IEEE Visual Communications and Image Processing (VCIP), pages 1–4. IEEE.

Cote, S., Butler, R., Michaud, V., Lavallee, E., Croteau, E., Mendrek, A., Lepage, J.-F., and Whittingstall, K. (2021). The regional effect of serum hormone levels on cerebral blood flow in healthy nonpregnant women. Human Brain Mapping, 42(17):5677–5688.

Cox, R. W. (1996). AFNI: software for analysis and visualization of functional magnetic resonance neuroimages. Comput. Biomed. Res., 29(3):162–173.

de Vos, V., Timmins, K., van der Schaaf, I., Ruigrok, Y., Velthuis, B., and Kuijf, H. J. (2021). Automatic cerebral vessel extraction in TOF-MRA using deep learning. In Landman, B. A. and Išgum, I., editors, Medical Imaging 2021: Image Processing, volume 11596, page 115962F. SPIE.

Forouhandehpour, R., Bernier, M., Gilbert, G., Butler, R., Whittingstall, K., and Van Houten, E. (2021). Cerebral stiffness changes during visual stimulation: Differential physiological mechanisms characterized by opposing mechanical effects. Neuroimage: Reports, 1(2):100014.

Gorgolewski, K. J., Esteban, O., Schaefer, G., Wandell, B. A., and Poldrack, R. A. (2017). Openneuro – a free online platform for sharing and analysis of neuroimaging data. Organization for human brain mapping.

Hartkamp, N. S., De Cocker, L. J., Helle, M., van Osch, M. J. P., Kappelle, L. J., Bokkers, R. P. H., and Hendrikse, J. (2013). In vivo visualization of the PICA perfusion territory with super-selective pseudo-continuous arterial spin labeling MRI. Neuroimage, 83:58–65.

Jiang, Y., Alaoui Mhamdi, M. A., and Butler, R. (2022). Inter-group heterogeneity of regional homogeneity (reho). bioRxiv.

Kalavathi, P. and Prasath, V. B. S. (2016). Methods on skull stripping of MRI head scan images-a review. J. Digit. Imaging, 29(3):365–379.

Livne, M., Rieger, J., Aydin, O. U., Taha, A. A., Akay, E. M., Kossen, T., Sobesky, J., Kelleher, J. D., Hildebrand, K., Frey, D., and Madai, V. I. (2019). A u-net deep learning framework for high performance vessel segmentation in patients with cerebrovascular disease. Front. Neurosci., 13:97.

Phellan, R., Peixinho, A., Falcão, A., and Forkert, N. D. (2017). Vascular segmentation in TOF MRA images of the brain using a deep convolutional neural network. In Lecture Notes in Computer Science, Lecture notes in computer science, pages 39–46. Springer International Publishing.

Silversmith, W. and Bae., J. (2020). Kimimaro: Skeletonize densely labeled 3d image segmentations. https://github.com/seung-lab/kimimaro.

Sun, Y. and Parker, D. L. (1999). Performance analysis of maximum intensity projection algorithm for display of MRA images. IEEE Trans. Med. Imaging, 18(12):1154–1169.

Sweeney, M. D., Sagare, A. P., and Zlokovic, B. V. (2018). Bloodbrain barrier breakdown in alzheimer disease and other neurodegenerative disorders. Nat. Rev. Neurol., 14(3):133.

Wang, R., Li, C., Wang, J., Wei, X., Li, Y., Zhu, Y., and Zhang, S. (2015). Threshold segmentation algorithm for automatic extraction of cerebral vessels from brain magnetic resonance angiography images. J. Neurosci. Methods, 241:30–36.

Xiancheng, W., Wei, L., Bingyi, M., He, J., Jiang, Z., Xu, W., Ji, Z., Hong, G., and Zhaomeng, S. (2018). Retina blood vessel segmentation using a u-net based convolutional neural network. Procedia Comput Sci, pages 1–11.

